# proDA: Probabilistic Dropout Analysis for Identifying Differentially Abundant Proteins in Label-Free Mass Spectrometry

**DOI:** 10.1101/661496

**Authors:** Constantin Ahlmann-Eltze, Simon Anders

**Affiliations:** Center for Molecular Biology, University of Heidelberg, Germany; Genome Biology Unit, European Laboratory for Molecular Biology (EMBL), Heidelberg, Germany

## Abstract

Protein mass spectrometry with label-free quantification (LFQ) is widely used for quantitative proteomics studies. Nevertheless, well-principled statistical inference procedures are still lacking, and most practitioners adopt methods from transcriptomics. These, however, cannot properly treat the principal complication of label-free proteomics, namely many non-randomly missing values.

We present *proDA*, a method to perform statistical tests for differential abundance of proteins. It models missing values in an intensity-dependent probabilistic manner. proDA is based on linear models and thus suitable for complex experimental designs, and boosts statistical power for small sample sizes by using variance moderation. We show that the currently widely used methods based on ad hoc imputation schemes can report excessive false positives, and that proDA not only overcomes this serious issue but also offers high sensitivity. Thus, proDA fills a crucial gap in the toolbox of quantitative proteomics.

## Introduction

Label-free quantification (LFQ) is a standard approach used in proteomics mass spectrometry (MS). Due to the similarity of this data type to expression microarray data, analysis methods from that field are commonly used for LFQ-MS. A major difference, however, is the presence of missing values in MS, but not in microarray data.

It is well established that missing values do not occur entirely at random, but more often at low intensities [1, 2, 3, 4]. The fraction of missing values varies by experimental design, but it is not uncommon to have more than 50% missing values, especially in affinity purification experiments. This issue hence cannot simply be ignored but needs proper handling, and doing so is a central challenge in statistical analysis of LFQ data, e.g., for identifying proteins which are differentially abundant between conditions. In the last years several method have been proposed to tackle this challenge, most of which rely on imputation, i.e., they simply replace missing values with some number that is deemed realistic.

However, a fundamental problem with imputation is that it obscures the amount of available information: imputed values will be considered as equally certain as actually measured values by any downstream processing (identifying differentially abundant proteins, clustering, quality control). This can invalidate inferential conclusions due to underestimating statistical uncertainty or cause loss of statistical power. Therefore, we propose a probabilistic dropout model that explicitly describes the available information about the missing values.

Figure 1A demonstrates that missingness carries information: observations in proteins with many missing values (red) have a lower intensity than observations in proteins with only one or no missing values (purple). In addition, Figure 1B illustrates that the ratio of these densities forms a curve with sigmoidal shape, clearly showing how the probability of a value being missing depends strongly on overall intensity.

**Figure 1:**
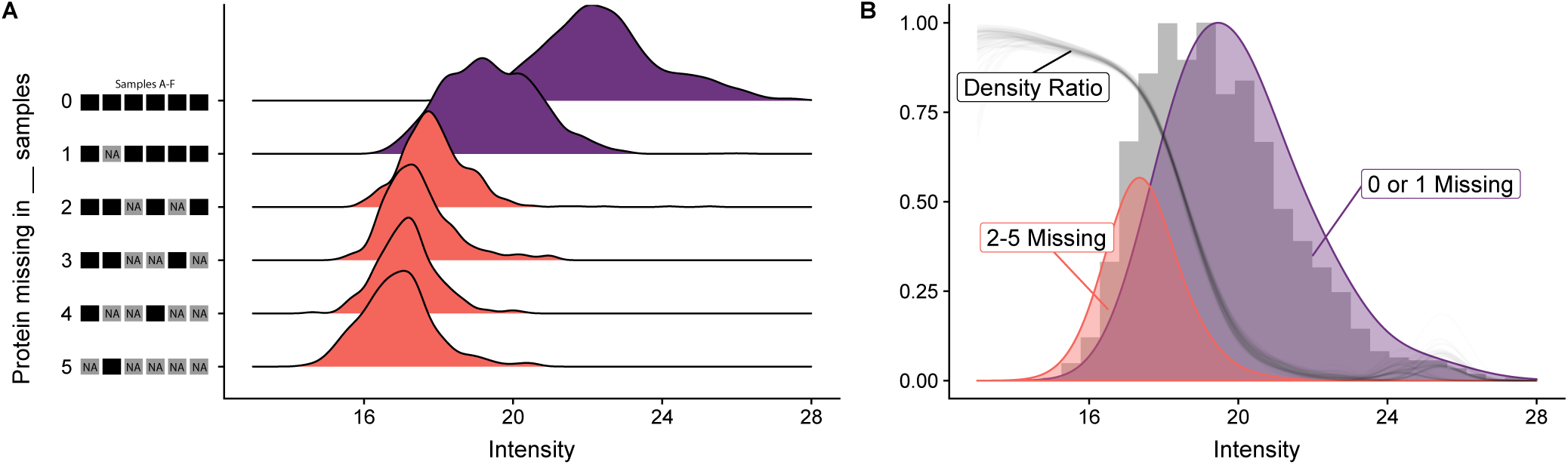
Missingness in label-free mass spectrometry is informative. (A) Intensity distribution for six replicates of the de Graaf data set. Shown is a ridgeline density plot of all the observed intensity values. They have been stratified by the number of samples in which the protein’s value was missing. The height of the individual densities is normalized per stratum. Panel (B) shows, in gray, a histogram of all the intensities. Overlayed are densities combining either the values from proteins with at most one missing value (purple) or with more then one missing value (red). The ratio of these two densities (gray line) has sigmoidal shape. The density ratio has been bootstrapped 100 times to show its sampling distribution.

If sample size is limited, substantial gains in statistical power can be gained from using shrinkage estimation procedure for variance estimation (“variance moderation”) [5]. This approach is widely used in transcriptomics data analysis, e.g., by the limma package [6]. The advantage of using limma or similar approaches for LFQ-MS has been advocated only rather recently (e.g., [7]). For example, the DEP package [8] performs imputation followed by a limma analysis to infer differentially abundant proteins. As stated above, the use of imputation may compromise the validity of limma’s statistical inference, and hence, the purpose of the present work is to adapt limma-style inference to account for values missing not at random and so improve power and reliability of differential abundance analysis for LFQ-MS.

A typical analysis of a label-free tandem mass spectrometry experiment consists of a number of steps. First, peaks in the MS1 need to be identified using the corresponding MS2 spectra. Second, the MS1 peaks need to be quantified. In the literature, two approaches are popular for this tasks: spectral counting and peak area integration [9]. Abundant peptides are more often recorded by the MS2, thus the number of MS2 spectra associated with a peptide can be used as a proxy for its abundance [10]. Alternatively, more abundant proteins cause larger peaks in the MS1, thus a second approach is to integrate the peak area of a peptide [11, 12]. Subsequent comparisons of the methods concluded that peak area based methods perform better than spectral counting [13, 14]. Consequently, we will only focus on methods that handle continuous intensities.

The third important step is the aggregation of the peptide level information to protein information. Traditionally, the peptide intensities are aggregated to protein intensities and then in a separated step the differential abundance is calculated for each protein. One popular method, that is directly integrated in the MaxQuant platform [15], is called MaxLFQ that uses delayed normalization based on the peptide ratios [16]. Alternative methods include summing up the peptide intensities, taking the average of the top three peptides [17], selecting a reference peptide to calculate the protein intensity [18], averaging the ratios [19], or using relative abundances while taking into account shared peptides [20]. More recently, some methods have been published that directly try to combine both steps to gain more power. The result of all those steps is a table with intensities for each protein and sample. The values in this table are commonly on a log_2_ scale in order to account for the mean-variance relationship of the raw data (Supplementary Figure S1).

Several methods have been published in the last years that use those protein intensities to calculate differential abundance. Perseus [21] is a platform with a graphical user interface, developed by the same group as MaxQuant, which provides functionality to normalize the data, impute missing values, identify significant proteins using a t-test and visualize the results. For multiple testing correction, Perseus offers two options: either the Benjamini-Hochberg procedure [22] or significance analysis of microarrays (SAM), a permutation-based correction originally described in Ref. [23]. As already mentioned, DEP [8] is an R package that provides a similar set of functionalities, but uses the more powerful variance moderated t-test to identify significant proteins using the R package limma [6, 24]. For multiple testing correction, DEP uses by default the methods in the fdrtool package [25]. In order to handle missing values, it provides an interface to a large number of imputation methods from the MSnbase R package [26]. In contrast, Perseus only provides two imputation methods, which either replace the missing values with a small deterministic value (MinDet) or with random values jittered around that small value (MinProb). DAPAR and ProStaR [27] are complementary software tools where DAPAR is an R-package that is similar to DEP, but has additional imputation methods based on the imp4p package [28]. ProStaR internally uses DAPAR and provides a web-based graphical user interface to make the software more approachable to newcomers.

Approaches that work without imputation are limited so far. one approach is the “empirical Bayesian random censoring threshold” (EBRCT) model, which avoids imputation by integrating over the inferred intensity distribution for missing values [29]. However, it cannot handle the case if a protein is completely missing in one condition. This can actually be problematic because in an affinity purification experiment those proteins might actually be the ones that we care about the most. Another tool that follows a similar idea is QPROT [4]; a command line tool that uses empirical Bayesian priors and integrates out the position of missing values using a cumulative normal distribution below a hard limit of detection.

Lastly, Triqler [31] is a tool that directly works on the peptide level instead of the protein level. This has the advantage that it can incorporate additional uncertainty due to the integration of multiple peptides to one protein intensity. It is a command line tool written in Python that fits an empirical Bayesian model integrating out the uncertainty for missing peptide quantifications.

Here, we present proDA (inference of *pro*tein *d*ifferential *a*bundance by *pro*babilistic *d*ropout *a*nalysis), a novel method for infering differential abundance that makes full use of the information inherent in missingness. proDA models the random process resulting in missingness and so avoids the problems discussed above that are inherent to imputation-based inference.

In the following section, we explain the intuition behind proDA and how it differs from the existing tools. In the third section, we use a spike-in and a semi-synthetic dataset to perform benchmarks. We show that many exiting methods have either low statistical power, or serious deficiancies in controlling false discovery rate (FDR), i.e., they refer too many false positives, and that hence the need for a general, powerful and statistically reliable inference method is unmet. We show that proDA offers strong performance with reliable FDR control, and thus is suitable to fill this crucial gap in existing methodology. In the fourth section, we demonstrate proDA in an application setting, by analyzing a real dataset studying ubiquitination. We close with a conclusion.

## Approach

The core of our idea is to combine the sigmoidal dropout curve for missing values with the information from the observed values. Figure 2 gives a conceptual overview of our approach. Our method works on the protein-level intensities, because this makes it compatible with a variety of different intensity aggregation methods, easier to use, and, as we will see later, the performance benefits from working on the peptide level are unclear.

**Figure 2:**
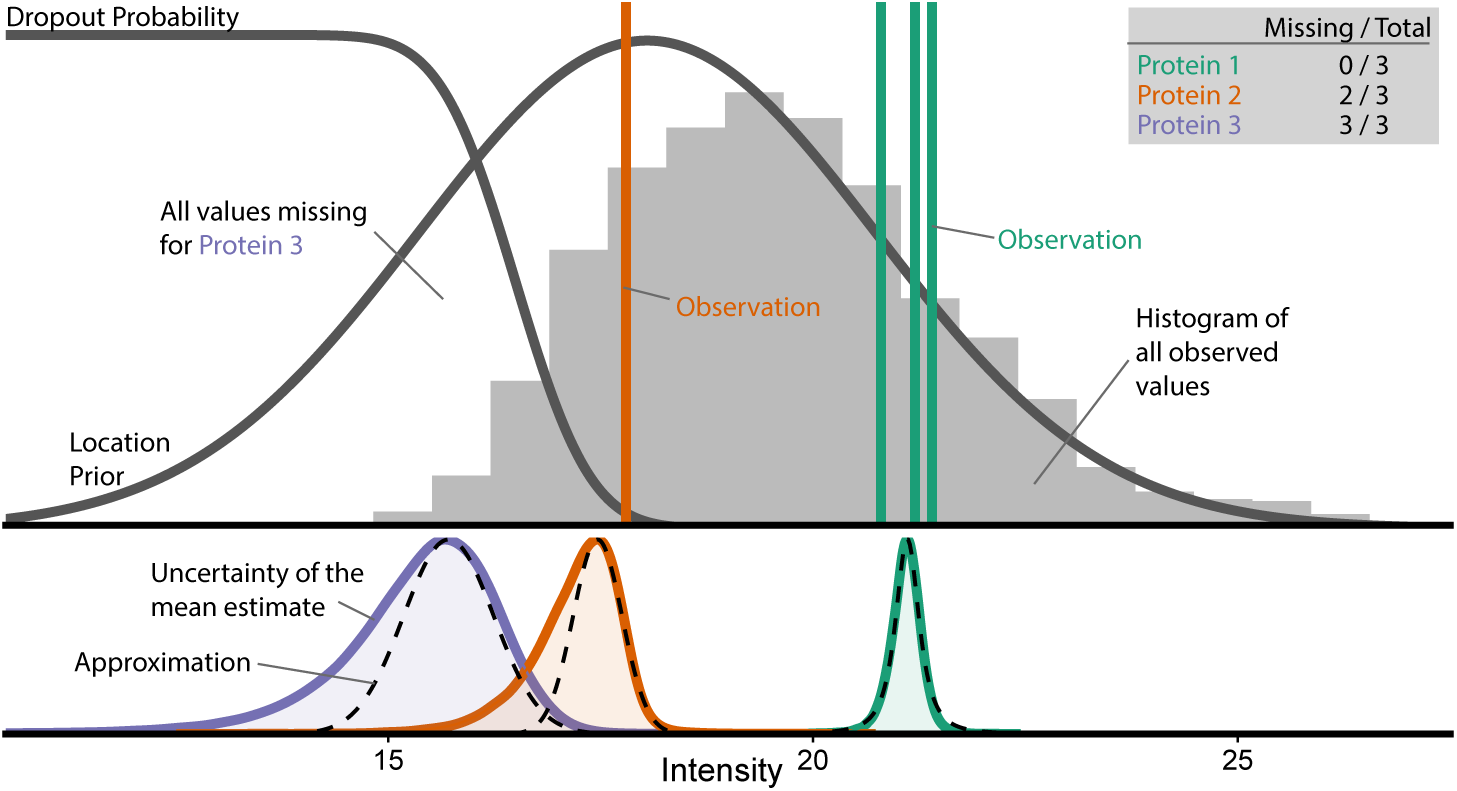
Intuition behind the probabilistic dropout model. We assume that the real intensity values approximately follow a normal distribution (“location prior”). The probability of not observing one of these values (“dropout probability”) is high for low intensity and low for high intensity. Hence, the distribution of the actually observed values (gray histogram) is skewed, with values missing in its left flank. The vertical lines indicate the observed intensities for three hypothetical proteins: Protein 1 (green) has complete observations, protein 2 (orange) has 2 missing values, and protein 3 (purple) has only missing values. The lower panel shows the inferred posterior probability distribution for the means for proteins 1, 2, and 3 (calculated using Stan [30]). The dashed lines show the symmetric approximation to these that we use for efficient inference.

All the mathematical details of our method are described in the Appendix, where we develop the approach for full linear models. Here, in the main text, we aim to provide a more intuitive explanation. We will first discuss the simple setting of an experiment with only a single condition with 3 replicates, and afterwards discuss inference of differential abundance between two conditions.

We assume that, within one condition, there is one expected value for each protein, the population average, i.e., the mean value we would get if we averaged over infinitely many replicate samples. The abundances in our 3 replicates scatter around this unknown “true” mean value, and our goal is to infer a posterior distribution that contains the true mean and captures our uncertainty about its location. In case of no missing values, such a posterior takes the shape of the t distribution, which is the basis for the well known Student’s t test (green posterior in Figure 2). Missing values cause these posteriors to become skewed, wider, and their mode (peak) to shift to the left of the average of the observed values (because the missing values are likely lower than the observed ones); see orange posterior in Figure 2. Even with no observed values, we can infer a posterior (purple posterior in the figure): its left flank follows the location prior, i.e., the distribution of values we actually expect in our data, and its right flank follows the dropout curve, because higher values would have likely been observed.

Hence, our approach first estimates from the data for all proteins the shape of the dropout probability curve and of the location prior. It then uses this information to infer for each protein an approximate posterior for its mean, with the necessary shift in mode location and widening due to the additional uncertainty from any missing values. We approximate the skewed posteriors with a symmetric approximation (dashed lines in the figure) that follows the right flank. (See Appendix, Section “Variance of the Coefficient Estimates” for details.)

The process of parameter estimation involves so-called shrinkage estimation (or moderation), which shares information across proteins in order to improve variance estimation (as originally proposed in Ref. [5] and also used in limma [6]). Furthermore, we apply shrinkage estimation not only to the variance but also to the location, as this enables us to handle the edge case of all observation missing in one condition.

To test if a protein is differentially abundant between two conditions, we compare the approximate posteriors inferred for the two conditions and calculate a p-value for the null hypothesis of both true means being equal. We do this using linear models, which allows for accommodating known covariates and complex experimental designs, in the same manner as limma offers for transcriptomics experiments.

## Validation and Comparison

We validated our approach and compared it with the existing methods discussed in the introduction. To this end, we used a dataset by de Graaf et al. [32], who analysed phosphorylation dynamics of Jurkat T cells over 6 time points using affinity purification. We only use the first time point, for which there are 18 samples, 3 biological replicates with 3 technical replicates each, which were measured in two separate mass spectrometry runs. As all these samples are in the same condition, we *a priori* do not expect any differences between the samples. We then introduce the changes ourselves so that we have a known ground truth. This has the additional advantage that we can vary the number of compared samples and the fraction of truly changed proteins to see how those affect the results. As a first test, however, we run our methods, as well as a variety of other methods, on the data as is, without any real differences between conditions, in order to check whether any of the tools might nevertheless falsely report statistical differences.

### Null comparison

Supplementary Figure S2 shows a heatmap of the data. There are many missing values (49%), which helps us to assess their impact on the different methods. In a typical affinity purification experiment, it is not unusual to have only three replicates per condition. So, we chose six samples and divided them into two synthetic conditions, ensuring that both contain a mix of different biological replicates, so that there is no signal in the null dataset (row marked “3v3” in Supplementary Figure S2).

We compare the 7 methods discussed in the introduction (Perseus, DEP, DAPAR, QPROT, EBRCT, Triqler, and proDA), running each tool with their default settings, except for the multiple testing correction, where we stick to the Benjamini-Hochberg method wherever possible, in order to make the results more comparable. DEP offers a range of different imputation methods; we chose to test it with five typical ones: Zero, MinDet, MinProb, KNN, and QRLIC. For DAPAR we used the structured least squares algorithm from the imp4p package [28, 33]. We ran QPROT with 2000 burn-in and 10,000 sampling iterations. For EBRCT, we have to filter out the proteins where for one condition all proteins are missing, because it cannot handle that edge case. Lastly, Triqler was a little more challenging to use, because it needs the data in a specific format that includes the decoy matches to calculate the FDR. For Triqler specifically, following the advice of the Triqler authors, we re-ran the MaxQuant quantification, with PSM and protein detection FDR set to 100% and matching between runs turned off, then converted the evidence file to Triqler input, skipping the normalization. We ran Triqler version 0.3.1 with the *fold change eval* setting changed to 0. It should be kept in mind that this means that Triqler was run on a slightly different data set without the same normalization and that thus the performance results are not one-to-one comparable.

Figure 3 shows the results: proDA correctly detects no significant proteins, as do most tested methods. The DEP method combined with zero imputation proves to be problematic, as are QPROT and EBRCT. DAPAR performance worse with increasing number of samples (“4v4” and “6v6” Supplementary Figure S3 and S4). Triqler (which is not included in the plots, because it does not calculate classical p-values) detects an unacceptably large number of false positives, which is why we will exclude it from the subsequent analyses.

**Figure 3:**
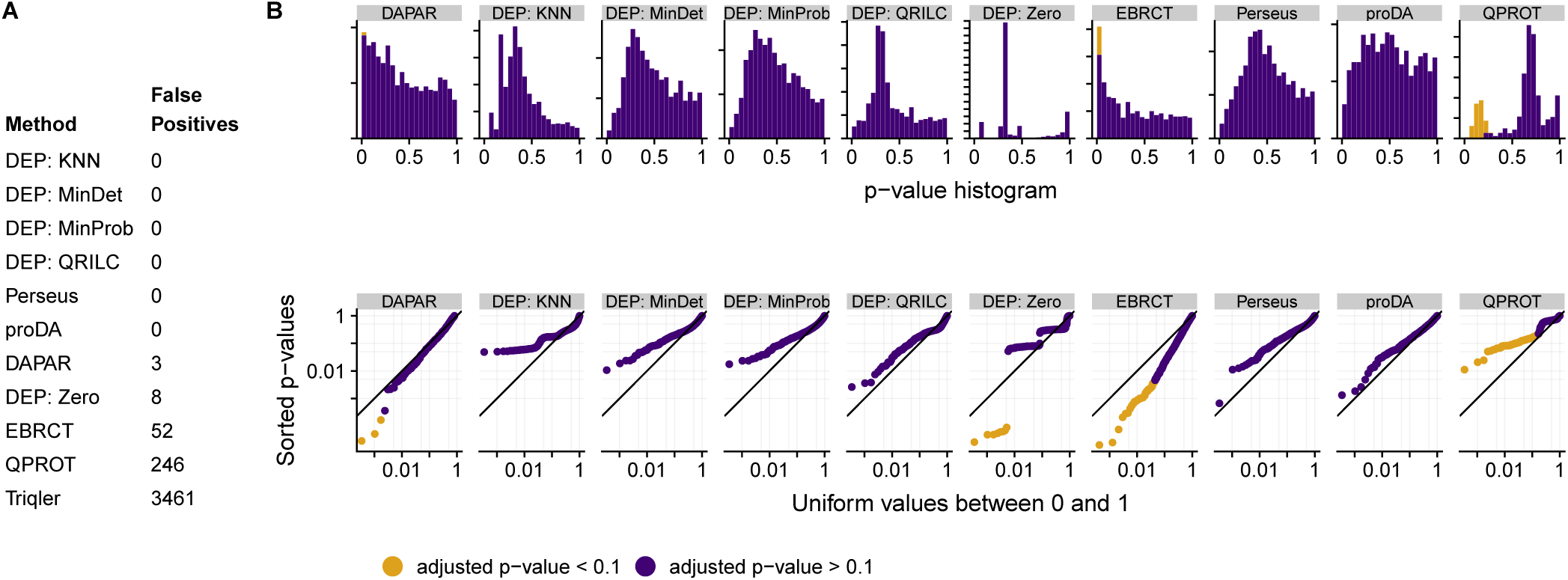
Comparison of the various tools’ ability to avoid false-positives: We have compared three replicates in the de Graaf dataset with three further replicates, i.e., with no differences, and would hence expect uniform p values and no detection of differentially expressed proteins after Benjamini-Hochberg adjustment. (A) Table of the number of falsely detected positives. (B) Histogram of p values. Ideally, these should be uniform. Each tick on the y-axis indicates 50 proteins. (C) Double-logarithmic QQ plots of p values; orange colour indicates proteins deemed significant at 10% FDR after BH adjustment. The QQ plots show the same data as the histograms but make it easier to judge uniformity for very small p values. Triqler is absent in the histogram and double QQ plot, because it does not calculate p-values, but directly calculates the FDR.

### Semi-synthetic dataset

We now introduce the artificial changes so that we have a known ground truth. We select 20% of all proteins and randomly shuffle those rows, but only in the first condition. This creates a realistic dataset where we know which proteins differ between condition one and two. The more common approach, where a selected number of proteins are shifted by a fixed effect size, is not applicable here, because shifting the mean of a protein would also imply a different probability for missing observations. Unfortunately, this idea can not be consistently applied to peptide level data, which is why we will have to leave Triqler out of the following benchmark.

We compared the other 6 method (Perseus, DEP, DAPAR, QPROT, EBRCT, and proDA), ran them with the same settings described in the previous section.

Figure 4A and B show the performance of the tools: in this test, DEP with most imputation methods (except zero imputation) and proDA succeed in controlling the FDR. EBRCT, QPROT, and DAPAR fail to control it. For those methods that passed the FDR control requirement, we can again ask which has most inferential power. Figure 4D shows the number of true positives that each method recovered depending on the desired FDR. proDA performs well in this test. Its actual FDR always stays below the desired FDR and at 10% desired FDR, it recovers 65% more true positives than the second best approach, DEP with MinDet imputation.

**Figure 4:**
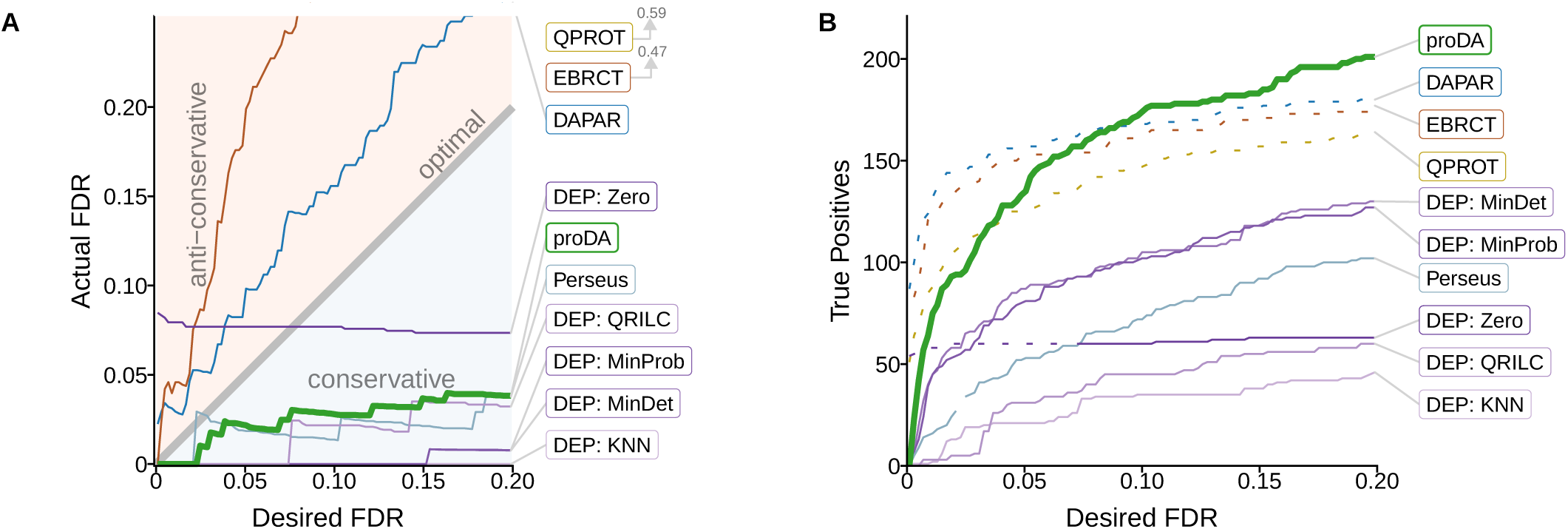
Performance comparison on the semi-synthetic data by de Graaf with three replicates and 20% changed proteins. A) Comparison of the user specified FDR (cut-off on BH-adjusted p value) with the FDR that is actually achieved by the tool according to the ground truth. (The line for QPROT is not shown as it is lierally off the scale.) B) Plot showing how many actually changed proteins (true positives) each method identified at a specified FDR level. The methods shown as dotted lines should be considered “disqualified” as they that failed to control the FDR in panel A.

The performance of DEP depended on the imputation method that is used. Zero imputation is problematic, as can be seen in this example, because it fails to control the FDR at small values. The best imputation methods are MinDet and MinProb, which perform nearly identical. Perseus with the MinProb imputation recovers fewer true positives than DEP, which is expected, because it uses the classical t-test and not the variance moderated version provided by limma.

In Supplementary Figure S5, we further distinguish the calibration and performance by the number of actually observed values in condition one and two. This shows that the failure of QPROT to control the FDR is because it has too many false positives specifically for proteins with zero against one observation. The opposite pattern is observed for EBRCT, that has too many false positive detections if a protein is observed in all conditions or mising in just one. proDA is more powerful than the other methods, because it shows consistently good performance across comparisons and in particular if only one or two observations are missing. DEP with the zero imputation method always identifies all proteins fully observed in one condition and completely missing in the other as significant. In many cases this is correct, but as this does not depend on the desired FDR, this can lead to an actual FDR that is too large if the user specifies a small FDR.

In Supplementary Figure S6-S8, we show that we mostly get consistent results even if we change the number of compared samples (3 vs 3, 4 vs 4, and 6 vs 6) and also if we change the percentage of true positive proteins (5%, 10%, 20%, and 30%). proDA reliably controls the FDR, except in two circumstances where the noise from the small number of changed genes (5% in Supplementary Figure S7A and Supplementary Figure S8A) causes a small violation of the FDR control. It also detects more true positives in most of the comparison than the other tools, especially whenever there are many missing values.

## Application

After demonstrating that only few tools, including proDA, control the FDR reliably, and that proDA is able to recover the largest number of changed proteins, we applied it to analyze a data set on the interaction landscape of ubiquitin [34]. In this example, we do not know the ground truth, but show that we recover proteins that biologically make sense. In the original publication the authors analyzed the data set using Perseus, later they presented the DEP R package for analyzing such data sets [8]. We will re-run the analysis that Zhang et al. describe with proDA.

Ubiquitin is a small protein that plays an important role in many different signaling pathways. There are three different kinds of ubiquitination: mono-ubiquitination, multi-mono-ubiquitination and poly-ubiquitination. Poly-ubiquitination is further distinguished by the linkage between the donor and the acceptor ubiquitin. The donor is linked with its C terminus to any of the seven lysines (K6, K11, K27, K29, K33, K48, K63) or the terminal methionine (M1) of the acceptor. Zhang et al. studied the recognition of those eight linkages and mono-ubiquitin by ubiquitin binding proteins. For this, they developed a new technique called ubiquitin interactor affinity enrichment-mass spectrometry (UbIA-MS) [8].

They run an enrichment experiment for each of the eight ubiquitin linkages plus one condition with mono-ubiquitin (Mono) and one empty control condition (ctrl). Each condition was measured in triplicates. To determine which proteins bind (directly or indirectly) to any of the ubiquitin linkages, we always compare the intensity for each protein to the corresponding intensity in the control group.

Figure 5 shows the results of the analysis with proDA. Figure 5A compares the total number of significant interactors at a nominal FDR of 10%, filtering out all proteins that had higher intensity in the control condition than in the ubiquitin condition. Figure 5B further stratifies the data from panel A. It not just describes how many proteins bind to a linkage, but also how many proteins bind to a specific combination of linkages. We can see that a majority interacts significantly with with all ubiquitins, but there are also proteins showing significant interactions only for specific linkages.

**Figure 5:**
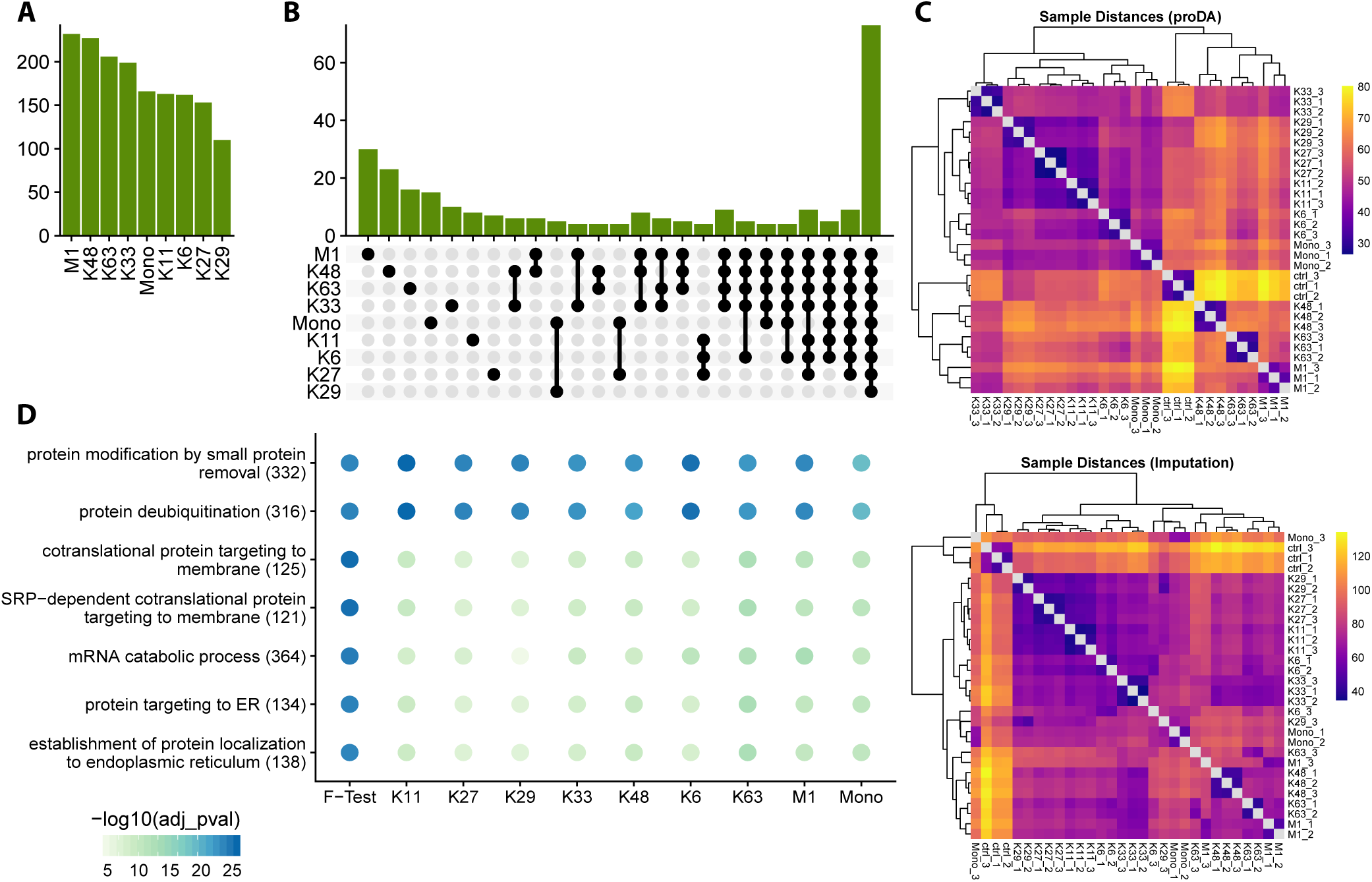
Ubiquitination analysis with proDA. A) shows the number of interactors for each of the nine conditions. The green bars show the number of proteins identified as significant interactors for each condition with proDA. B) breaks down those interactors into more detail. It shows the number of proteins that interact with a specific combination of ubiquitin linkages. The total number of intersections was limited to the largest 25 sets ordered by degree. C) shows two heatmaps with the sample distances, calculated according to Equation (42) (upper heatmap) and on the imputed dataset (lower heatmap). The rows and columns were clustered using hierarchical clustering on the distances. D) shows a dot plot with the seven most significant gene ontology (GO) terms related to the set of interactors with any of the nine conditions and the set of proteins that differ over all conditions (“F-test”).

Figure 5D demonstrates that proDA has not just recovered many interactors, but proteins related to gene ontology sets relevant for ubiquitination [35, 36, 37]. In addition to the 9 ubiquitination conditions, here we also list the results of conducting an F test to identify all proteins that differ in any condition, as an example for the ability of proDA to perform missing value aware ANOVA.

## Distances

A commonly used approach for sample quality control is to calculate some measure of similarity for all pairs of samples, in order to check whether replicate samples appear more similar than samples from different conditions.

Typically, Euclidean distance is used, e.g. in the ubiquitilation study by Zhang et al. [8] who use MinProb imputation before Euclidean distance is calculated. Figure 5C shows the outcome of this procedure for the ubiquitin data. Differences in the shape of the dropout probability curve can strongly influence a distance calculated in this manner. Based on the proDA model, we developed an approach to calculate Euclidean distance in a probabilistic manner without the need for imputation in order to reduce the effect of differences in dropout probabilities (Appendix, section “Distances”)). In fact, our distance calculation is able to recover the triplet structure of the data set, while the MinProb imputation based distances do not (Figure 5C).

## Conclusion

In this paper, we have presented our R package proDA for identifying proteins that are differentially abundant in label-free mass spectrometry data sets. The main challenge for analyzing label-free mass spectrometry data are the large number of missing values. We suggest to handle them using a probabilistic dropout model combined with empirical Bayesian priors to combine the available information from observed and missing values.

In the performance comparison with existing tools with and without any true positives, we saw that some method produce a lot of false positives. In particular, Triqler had difficulties on the null-comparison. This could be due to the challenges of balancing the different levels of uncertainty from the missing values and the observations on the peptide level.

On the semi-synthetic data set, we saw that proDA recovers more true positives, while controlling the false discovery rate. We showed that imputation can be problematic because it either leads to a loss of power or worse to not controlling the false discovery rate. The improved sensitivity of proDA comes at the prize of a somewhat increased run time. Whereas DEP, DAPAR, and Perseus finish within seconds, our model might need one or two minutes to calculate a result. In the end, we believe the increased computational demands are justified, because the analysis run time is still fast enough for interactive use.

In addition, our tool can handle any design specified as a linear model. This has the advantage that one can not just fit two condition comparisons, but also time series data, nested data with patient and treatment specific effects, and account for known covariates in the model. Besides proDA, this is only supported by DEP although this is standard for transcriptomic tools.

In conclusion, we have demonstrated that imputation can be problematic and that properly modelling the uncertainty posed by missing values boosts power.

## Availability

### Software

The proDA method is implemented as an open-source R package. Documentation, installation instructions and download links are provided at the BioConductor package repository https://www.bioconductor.org/packages/proDA/.

### License

proDA is made available as open-source software under the GNU General Public License, version 3 or later.

### Source code for software

Full source code for the software is available on GitHub, at https://github.com/const-ae/proDA

### Source code for benchmarks and example analyses

The full R code to produce all the figures and all the benchmark and example application results reported here is available at github.com/const-ae/proDA-Paper as R notebooks. The example data used was downloaded using the accessions given in Refs. [32] and [34]; details are given in the R notebooks.

## Appendices

### Mathematical Description of the Probabilistic Dropout Model

At the center of proDA is the idea of a probabilistic dropout model. As we saw in Figure 1, the chance of a missing value decreases with increasing protein intensity. We will model this relationship with a sample specific sigmoid dropout curve. We will describe our model as a generative model, i.e., the mathematical relations that we believe could be responsible for the data table from which we start our analysis.

We denote the data table as a matrix *Y* with *I* × *J* rows and columns, where *I* is the number of proteins and *J* is the number of samples, and the matrix elements *y*_*ij*_ are the recorded MS1 intensities (on the logarithmic scale) for protein *i* in sample *j*. We write *y*_*ij*_ = `NA` if the intensity value for protein *i* is missing in sample *j*.

As usual in linear models, we describe the experimental design with a design matrix *X* with dimensions *J* × *p*. For the expected intensity value for protein *i* in sample *j*, we write *µ*_*ij*_ = *X*_*j*_***β***_*i*_, where *X*_*j*_ is the row of the design matrix *X* corresponding to sample *j*. Our goal is to find for each protein *i* the *p* coefficients of the vector ***β***_*i*_.

We assume that the actual log intensities *z*_*ij*_ scatter around their expected values *µ*_*ij*_ according to a normal distribution with variance 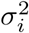. As discussed in the main text, the values *z*_*ij*_ are not always recorded; with some probability, they might suffer a drop-out. If they are observed, we set *y*_*ij*_ = *z*_*ij*_, otherwise we set *y*_*ij*_ = `NA`. We assume that the probability of a dropout depends on the intensity *z*_*ij*_ and we will model this with a sigmoidal relationship. There are several possible functions describing curves with sigmoidal shape; for mathematical convenience, we chose the inverse probit, i.e., the complement of the cumulative density function (CDF) of a Normal distribution. In formal notation, this model is

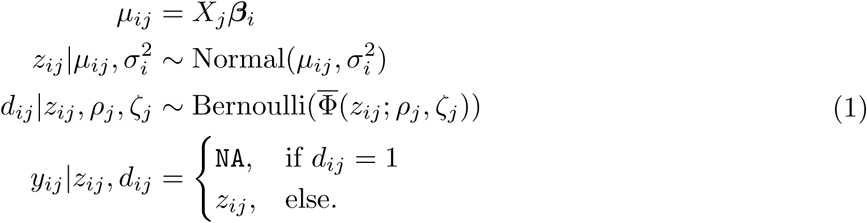

Here, *z*_*ij*_ are the latent intensities, that we do not have full access to because of the dropouts. ***β***_*i*_ are the coefficients for which we want to find out if they or their linear combination are different from zero. *d*_*ij*_ indicates if a protein is missing in the specific sample. The probability of missingness (*d*_*ij*_ = 1) is given by the sigmoidal dropout curve 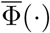. As already mentioned, we chose to describe this sigmoid with the complement 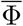 of a Normal CDF Φ, parameterized using the inflection point *ρ*_*j*_ and the scale *ζ*_*j*_:

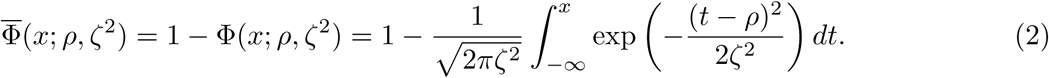

The decreasing behavior of 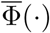 matches the observation that measurements with lower intensities are more likely to be missing.

In addition, we assume that the means *µ*_*ij*_ and the variances 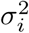 are similar across proteins and add the priors

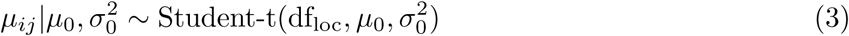

and

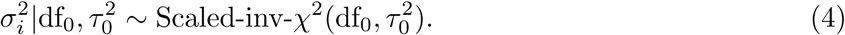

The prior in Equation (3) on the protein means *µ*_*ij*_ is important to handle the edge case if in one condition a protein is completely missing. The prior in Equation (4) corresponds to the variance moderation of limma [6].

The probability density function of the generalized Student’s t-distribution used in Equation (3) is defined as

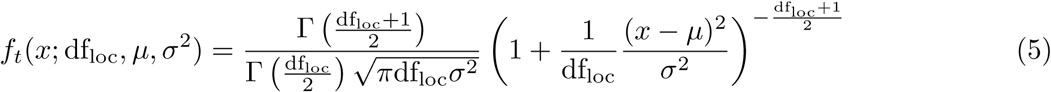

and the probability density function of the scaled inverse *χ*^2^ distribution used in Equation (4) is

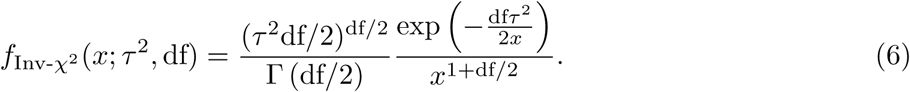

We iteratively estimate the hyper parameters and the protein specific parameters using a maximum *a posteriori* approach until the model converges. To identify which coefficients in ***β***_*i*_ are significant, we use a Wald test or likelihood ratio F-test [38].

### Model Fitting

In the following section, we will explain how to infer the feature parameters ***β***_*i*_ and 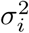, and then the hyper-parameters 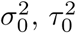, df_0_, ***ρ*** and ***ζ***. We assume that df_loc_ is fixed by the user.

To simplify the notation, we will first focus on only one protein and assume that all samples belong to the same condition and thus suppress all subscripts *i* and *j*. This also allows us to directly talk about *µ* instead of *X****β***, because in that specific case they are identical.

If there were no missing values and if we ignored the priors, the likelihood of *µ* and *σ*^2^ given the observations ***y*** would be

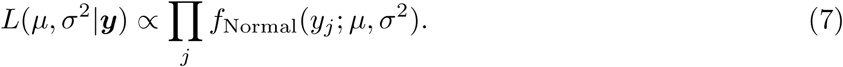

To handle the mix of observed and missing values in ***y***, we will extend the above equation by marginalizing out the missing values

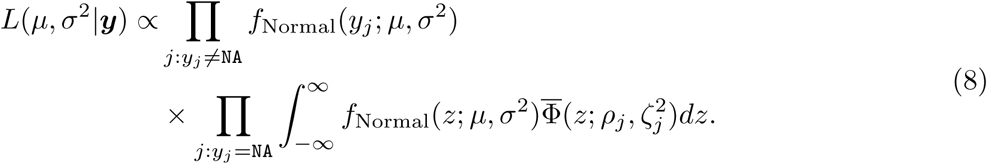

The integral in Equation (8) can be simplified

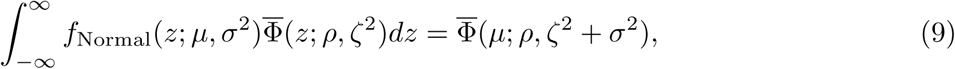

with the proof provided, for example, in Refs. [39] and [40].

Now, we can combine Equation (8) and Equation (9), add the priors that we proposed in Equation (3) and Equation (4) and use *X****β*** instead of *µ*. We find that the joint density is

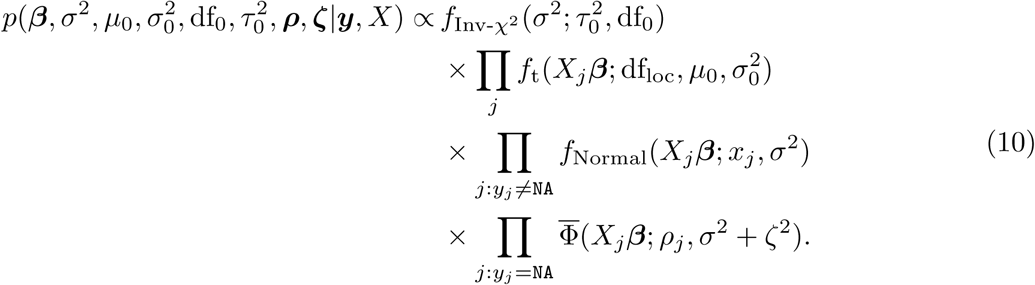

In this likelihood, we have written the hyper-parameters (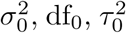, ***ρ*** and ***ζ***) as known constants. In practice, we use an iterative procedure, alternating between two steps: (i) We first fix the hyper-parameters to estimate the feature parameters and their uncertainties in a Bayesian fashion. We calculate the first two moments of their posterior associated with Eq. (10). (ii) Then, we fix the feature parameters to estimate the hyper-parameters by maximizing the likelihood (10). We iterate between those two steps until the estimates have converged. In the following sections, we explain either step in turn.

One might argue that the lack of proper hyper-priors renders our approach not a “pure” Bayesian one. Alternatively, however, one may read Eq. (10) as implicitly including improper uniform hyperpriors for the hyperparameters. This is permissible because in an empirical Bayes approach, the hyperparameters are estimated from a a very large amount of data (here, from sharing information over all proteins), and hence the likelihood associated with this empirical data so strongly dominates the hyperpriors that the specific choice of hyperpriors is negligible. This also leads to the posteriors for the hyperparameters being very narrow, which justifies our approach of treating the hyperparameters as constants in step (i) and performing step (ii) by simply maximizing the likelihood. We note that the same approach is taken in other empirical-Bayes approaches (e.g., in limma [6]) even though this fact is often not stated explicitly.

### Feature Parameter Estimation

For each feature (protein), we search the linear model coefficient vector ***β***_*i*_ and the corresponding variance value 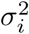 that best explain the observed intensity values **y**_*i*_ in the sense that they have high a posteriori probability according to Eq. (10). We approximate the posterior for (***β***_*i*_, *σ*_*i*_) as a multivariate Gaussian, i.e., we seek the first two moments of the posterior.

To this end, we start by using a maximum *a posteriori* (MAP) approach to find the 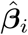 and 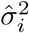 that maximize Eq. (10) for the observed intensity values **y**_*i*_ and a given set of hyperparameter values:

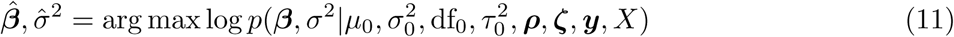

with

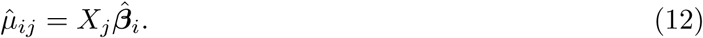

In proDA, the function `pd_lm` performs this optimization for each gene *i*: We take the logarithm of Equation (10) and derive its Jacobian and Hessian to efficiently find the mode. To this end, we use the `nlminb` function in R that wraps the PORT routines [41] for the actual optimization.

Then, `pd_lm` continues with the following steps.

#### Coefficient Estimates

For the posterior of the coefficients vector ***β***_*i*_, we simply use a multivariate Gaussien with the MAP estimate 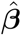 as mean. For the posterior’s variance-covariance matrix, see below.

#### Unbiased Variance Estimates

For the coefficients, the MAP estimates 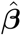 can be used, but for 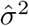, we have to take its bias into account.

In a standard linear model, we would expect the maximum likelihood estimator 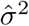 to be biased and underestimate the true variance *σ*^2^ by

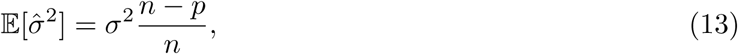

and hence correct 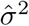 by multiplying it with *n/*(*n* − *p*).

With missing values, the challenge is that it is unclear what should be used for *n*. The simple approaches of setting *n* = *J* or *n* = |{*y*_*j*_ ≠ `NA`}| are problematic because they over- or underestimate the amount of information from the missing values. Instead, we will estimate an effictive value for *n* from the variance of the 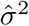 estimate at the mode, which is given by

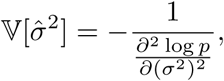

where *p* is the posterior given in Equation (10). We get this second derivative for free as an element of the Hessian matrix ***H***, which is calculated anyway during the maximization of log *p*:

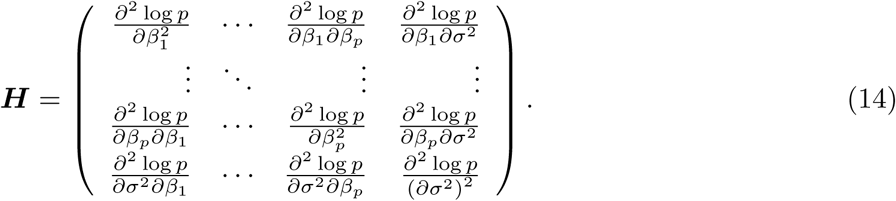

We find the value of *n* using an analogy to the standard linear model without missing values. If we have some values *y* and use their mean 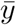 as the mean estimate 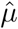, the density of *σ*^2^ would be

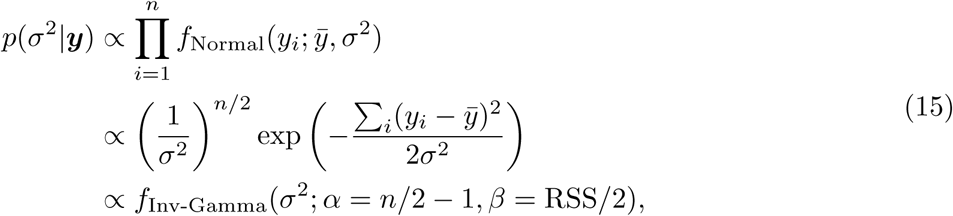

where 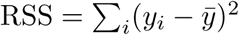. The mode of the inverse gamma distribution is

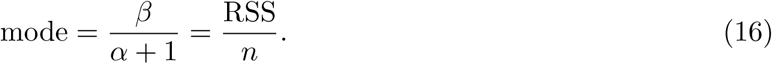

We can now find the expected value of the second derivative that the inverse gamma distribution has at the mode, which is

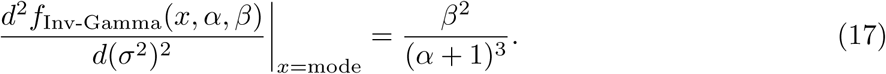

With missing values, we can still identify identify the MAP for 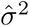 and the associated uncertainty 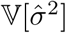. If we now plug in those values

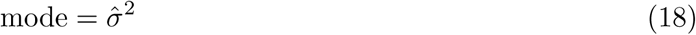

and

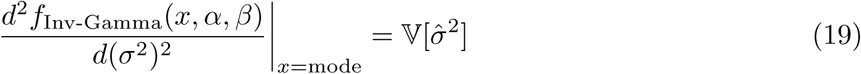

and solve Equation (16) and (17) using Equation (15) for *n* and RSS, we find that

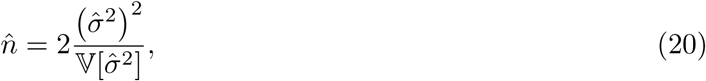

and

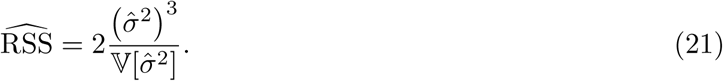

Finally, we can now identify the unbiased estimate of the variance, which is

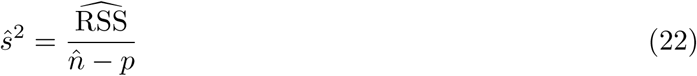

and estimate the degrees of freedom

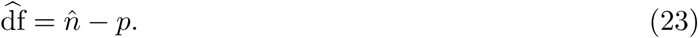

Sometimes 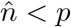, in which case we fix 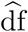 to a small, but positive value (ie. 0.001), and estimate

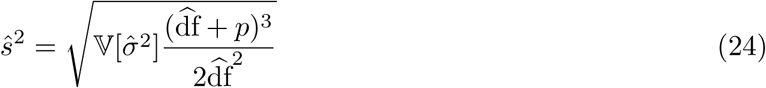

so that the approximation matches the scale of the original distribution, although the mode is slightly off.

#### Variance of the Coefficient Estimates

In a standard linear model, it is easy to find the standard error for the coefficients because it is simply

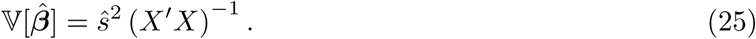

Again, this cannot be directly applied to the case with missing values, because it is possible that 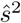 is small, but we nevertheless are very uncertain of *β*_*j*_ because there are many missing values for that coefficient. Instead, we will use the inverse of the Hessian of the log likelihood with respect to the coefficients, which we calculate using the unbiased estimate 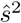

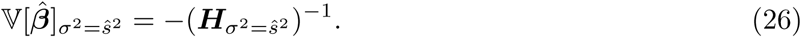

This works well for the cases where the distribution of *β*_*j*_ does not have too much skew. However, in Figure 2, we saw that for the cases with many missing values the skew can be considerable. If we just used the curvature (Hessian) at the mode, we would be wrong on both sides of the distribution. On the left, the approximation would be too narrow and on the right, it would be too wide.

We know that the distributions with considerable skew are always on the low end of the intensity distribution. In the typical comparison, we care about the inside flanks of the two distributions that we compare. We know that the skew will be larger for the distribution with the lower intensity. To minimize the approximation error, it is thus more important to correctly model the right flank of the distribution. Otherwise we would unnecessarily lose power. In the performance evaluation (Section “Results”), we can see that this approximation and symmetrization seems permissible and does not unduly inflate the false discovery rate.

We will hence calculate a correction factor, with which we scale the Hessian-derived variance, that reduces the variance such that we match the right flank of the distribution. If there is no skew, we know that if we go *k* units from the mode 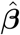 in the direction of *β*_*i*_ the log probability should decrease by 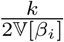, because the log density should behave like a multivariate parabola. Note that we still use the Hessian with 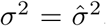. In our implementation we use *k* = 8 which ensures that the tails, which we care about the most, are approximated best. Figure 6 shows an example for the one dimensional case.

**Figure 6:**
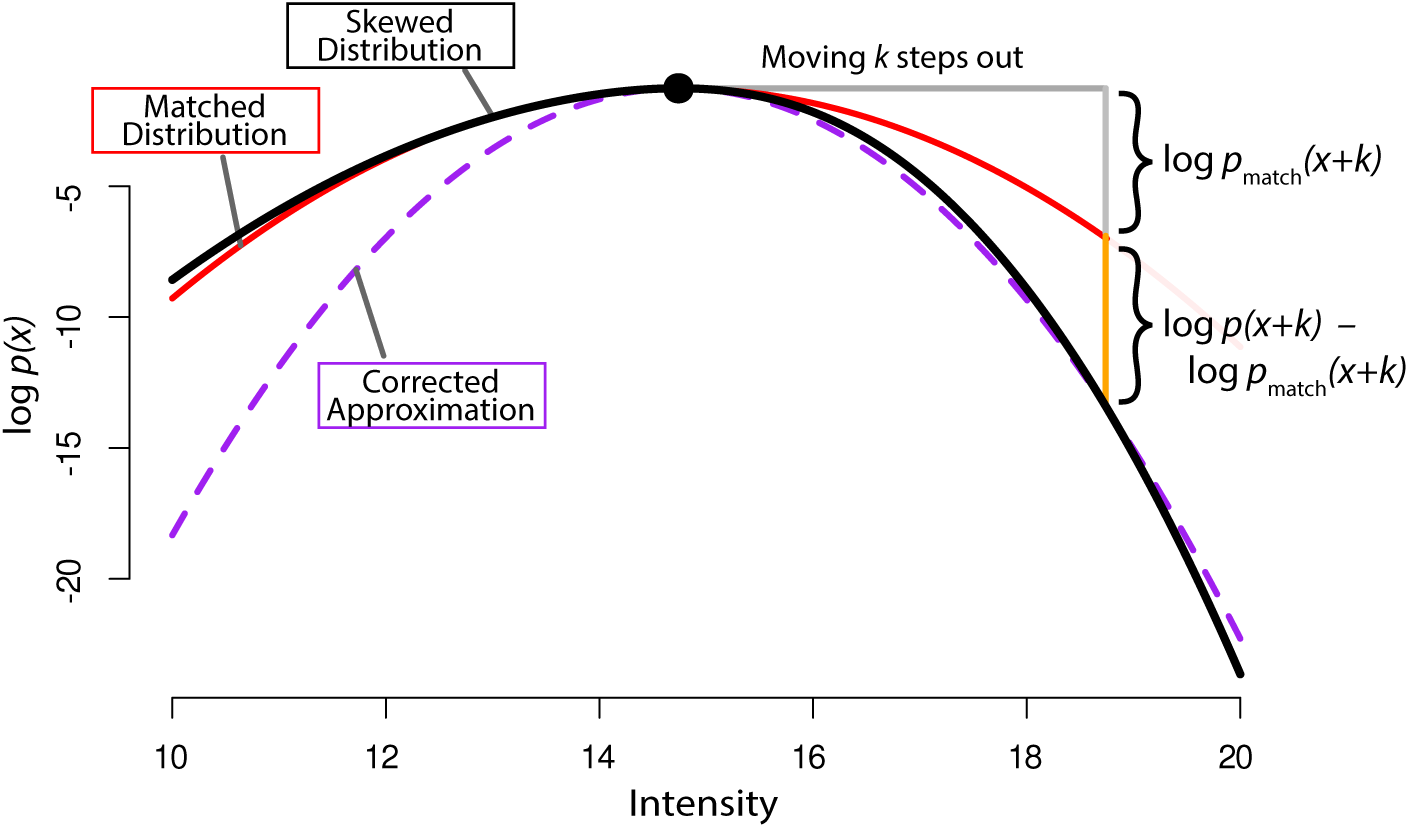
Sketch of how the correction factor is calculated and how it modifies the approximation in the one dimensional case. The original skewed distribution is shown in black. The black circle marks its mode. The red parabola (which is equivalent to a normal approximation directly using the Hessian) has the same curvature at the mode but is too wide on the right-hand side. If we move out *k* steps from the mode to the right and calculate how much more the actual density has decreased than we would expect from the Hessian (orange section), we can calculate the correction factor and construct an approximation that captures the behaviour of the right flank better (purple parabola).

From this relation we can calculate the correction factor which is

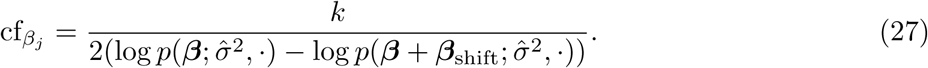

Here, we cannot directly use 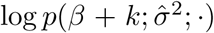, because in the multi-dimensional case we have to correct the distance we go out by the changed width of a multivariate Gaussian. Instead we use ***β***_shift_ which is a vector of zeros, except for the *j*’s entry which is

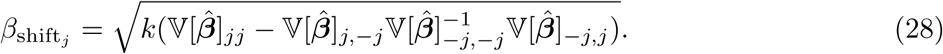

Here we use the notation 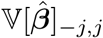 to mean that we take the *j*-th column of 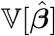 and remove the *j*-th entry. Similarly, 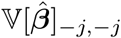 means that we remove the *j*-th row and column from 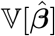. To understand Equation (28), note the similarities to the variance of the conditional distribution along an axis of a multivariate Gaussian.

We then identify the final covariance matrix as

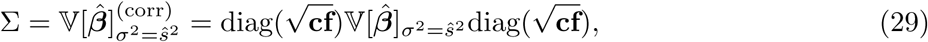

where **cf** is the vector formed by the correction factors from Equation (27).

This covariance matrix is then used to perform the Wald tests for inference of statistical significance of coefficients or contrasts. For a given contrast vector ***c***, we get a *t* statistic

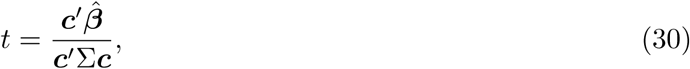

for which we report the two-sided tail probability of a t distribution with *I* − *p* degrees of freedom as the p-value. In proDA this is implemented in the `test_diff` function.

### Hyper-parameter Estimation

In the previous section, we have focused on individual proteins and suppressed the subscript *i* and handled ***y*** as a vector of size *j*. Now, we will describe how to fit the hyper-parameters across proteins and thus mention *i* and work with the full data matrix *Y*.

We estimate the hyper-parameters using an empirical Bayesian approach. Unlike a pure Bayesian setting where we would have hyper-priors on those parameters, we rely on the observed data to get accurate point estimates of the parameters for the dropout curves, the variance prior, and the location prior. This is typical for the empirical Bayesian framework, because the parameters are estimated across all proteins which means that the residual uncertainty of the hyper-parameter estimates is very small compared to the estimates of the feature parameters.

#### Dropout Curves

We fit one dropout curve for each sample, because the number of missing proteins can differ substantially between samples and the effect cannot be fixed by normalization. The drop-out probability for sample *j*, given in Equation (2), is parametrized by *ρ*_*j*_ and *ζ*_*j*_. We find these by maximizing the following log likelihood

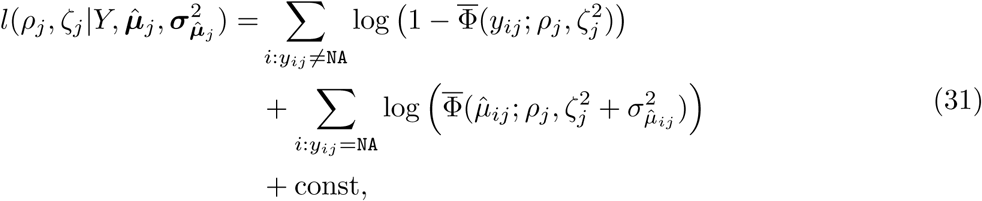

where we have plugged in the predicted values 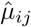 for the missing observations and the associated uncertainty

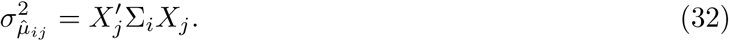

We use the general purpose optimizer implemented by the R function `optim` [42, 43, 44, 45] to find the maximum. The code performing the described step can be found in proDA’s internal function `dropout_curve`.

#### Variance Prior

In the model without missing values, Smyth [6] has described how to estimate the hyper-parameters of the variance prior 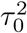 and df_0_ from the unbiased variances 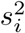 and the degrees of freedom df_*i*_. He showed that the unbiased variance estimates 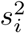 follow a scaled F distribution:

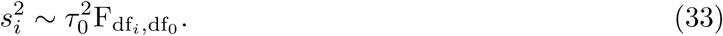

The scaled F distribution has the density

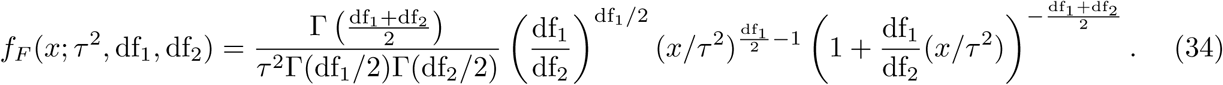

Smyth [6] provides closed form estimators for 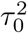 and df_0_ based on the moments of the logarithmized distribution, which are approximations of the maximum likelihood solution for Equation (34).

In the previous section (Equation (22) and (23)), we have shown how to find 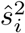 and 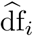 in the case of missing values. However, if we were to use these values directly we would have the problem that they already contain the information of last round’s hyper-parameters, and thus the variance prior would get narrower and narrower. To avoid this problem, we recalculate the quantities from the last section without location and variance moderation (simply by ignoring the first two lines of Equation (10)) and call them _*u*_*ŝ*^2^ and 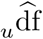. Here, *u* stands for un-regularized. We use those values for the inference of 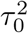 and df_0_, which are just the quantities that maximize the log likelihood of the scaled F distribution:

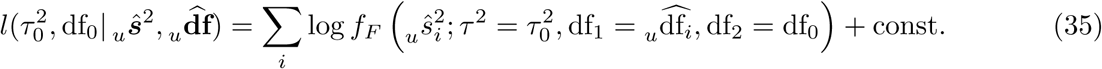

In proDA, the variance prior estimation is performed by the internal function `variance_prior`.

##### Location Prior

Lastly, we explain how to find the hyper-parameters for the prior on the protein means. Equation (3) states that we believe that the proteins means *µ*_*i*_ are drawn from a Normal distribution. We estimate the mean of that location prior using a trimmed mean of the predicted values across all proteins and samples

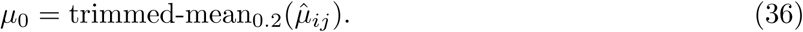

However, we cannot calculate the variance the same way, because using the already regularized values 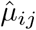 would lead to narrower and narrower estimates. This means that we need the un-regularized value 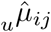.

We are only able to calculate 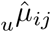 if we have at least one observation, but it is more likely to have proteins without any observations left of the global mean *µ*_0_. Thus, we will ignore all 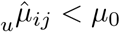 and assume that the distribution is symmetric.

To find the empirical Bayesian estimate of 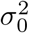, we use the approach described in Ref. [46] for a Normal prior density, which shows that 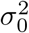 is the value that solves

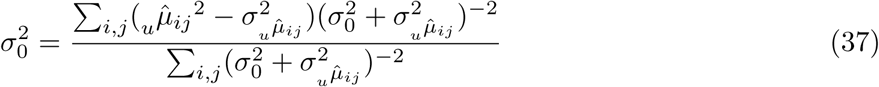

which we find using the `root` function in R.

This step is performed by the internal function `location_prior`.

### Distances

Understanding which samples are similar and which are not is an important step for quality control. Again, missing values make this task difficult: we cannot use the log intensity values for all proteins to form one vector per sample and calculate Eudlidean distances between these vectors. If we were to impute the missing values, we would get unrealistically high similarity for samples with many missing values. Instead, we propose to construct a probabilistic similarity measure.

The most commonly used measure of sample similarity is the Euclidean distance between two samples in the feature space. If there were no missing values this would just be

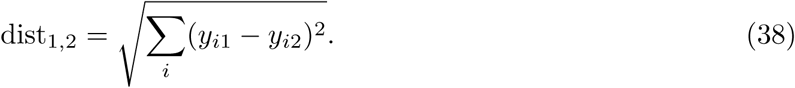

The feature space is the *I* dimensional space where each axis corresponds to one protein and each sample is a point ***x***_*·j*_ in that space. If a protein measurement is missing, we know that its intensity was low, but we cannot exactly say where along that particular axis the point is. We therefore consider the coordinates of ***x***_*·j*_ as normally distributed random variables, 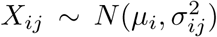. For observed intensity values *y*_*ij*_, we set *µ*_*ij*_ = *y*_*ij*_ and *σ*_*ij*_ = 0. For missing values, we set *µ*_*ij*_ to our estimate according to Equation (12) and 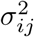 to the standard error of this estimate according to Equation (32). Then, the distance between two samples (here labelled 1 and 2) becomes a random variable, too, defined as

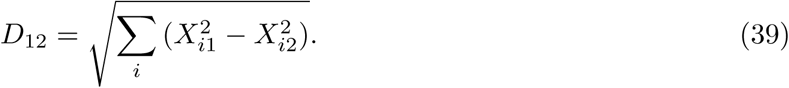

The full distribution of *D*_12_ may be hard to derive, but Ref. [47, p.53] provides formulas to analytically calculate the moments of the squared distance 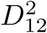:

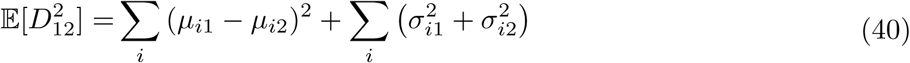

and

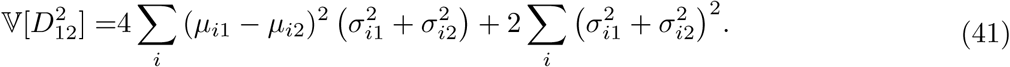

We can use those equations to approximate the actual quantities of interested: the estimated distance and the associated uncertainty

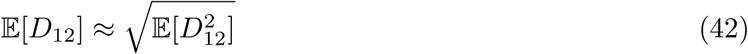

and

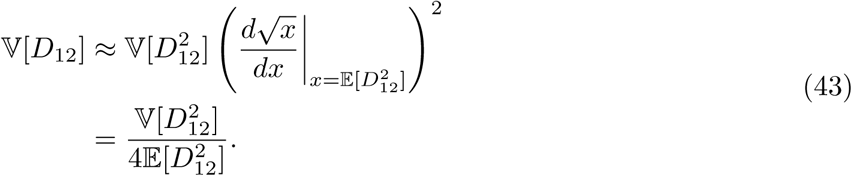

The proDA sample distance heatmap shown in Figure 5C are the expectations from Equation (42), calculated with proDA function `dist_approx`.

## Supplementary Figures

**Suppl. Figure S1:**
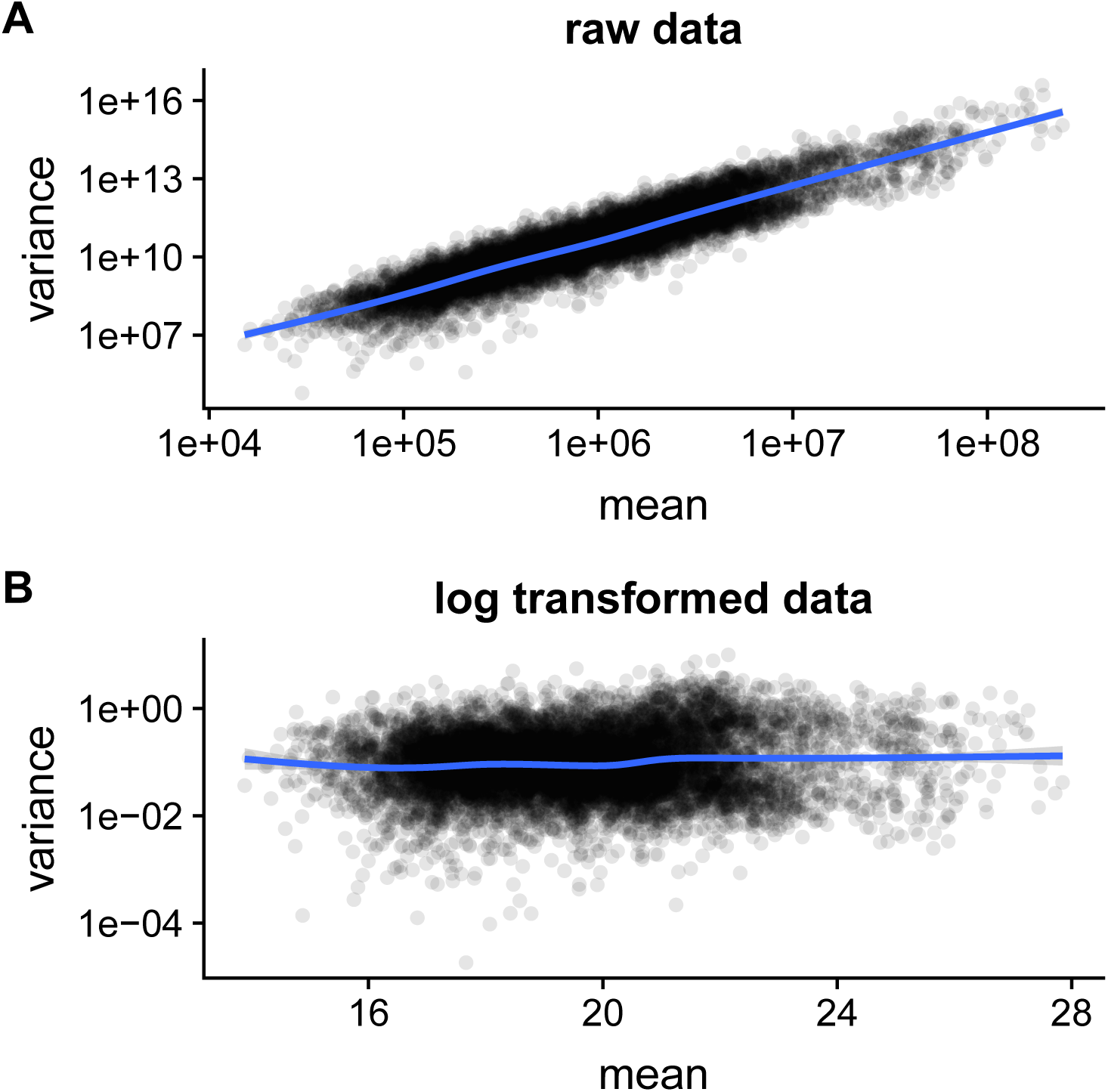
Mean-variance relationship on the full de Graaf dataset. Each dot represents mean and variance for one protein at one time point and MS run. The blue line is a ggplot2 smoothing fit. A) the mean-variance relation on the raw data. B) the mean variance relation on the log_2_ transformed data.

**Suppl. Figure S2:**
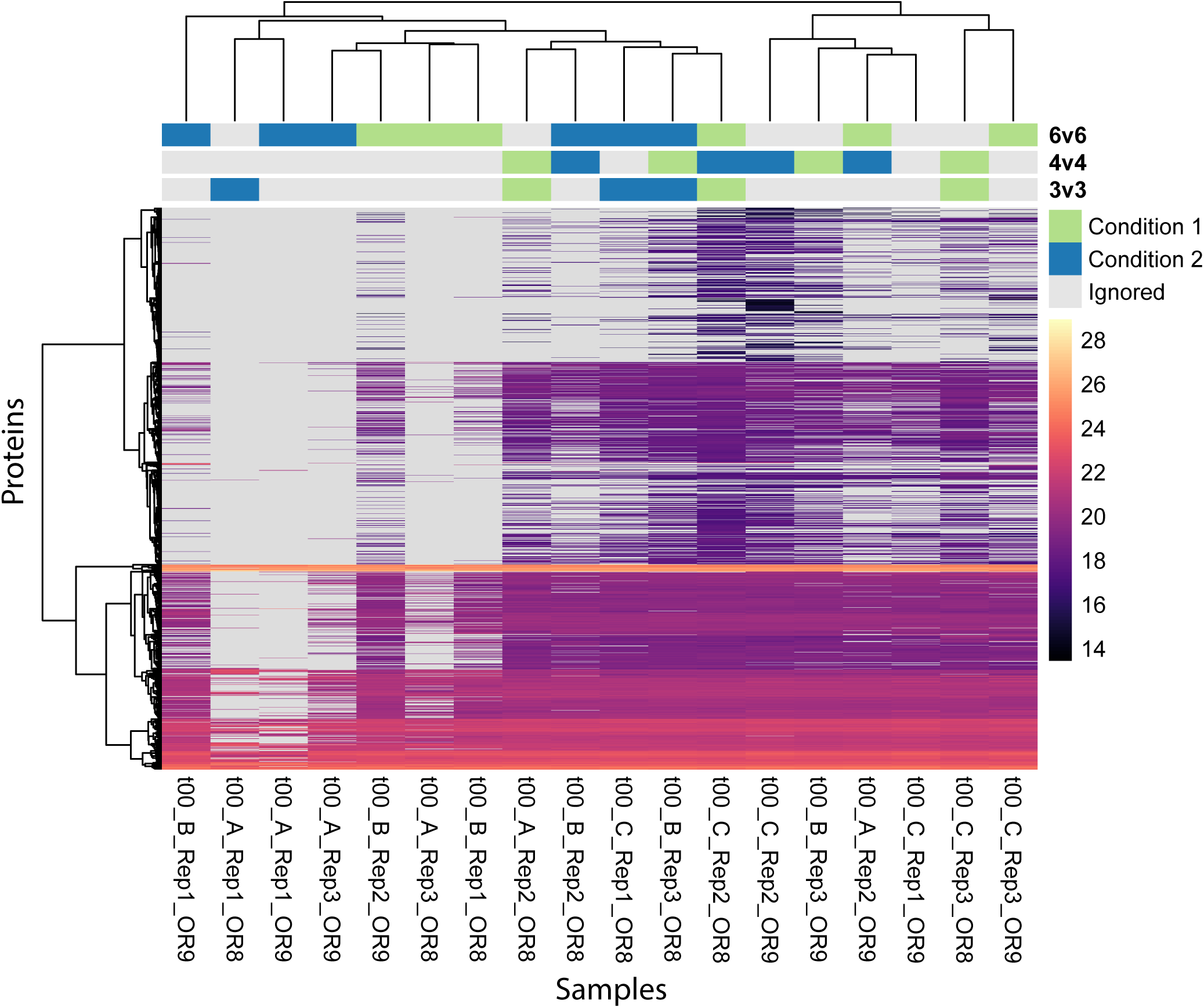
Heatmap of the de Graaf dataset. Each column is a sample and each row is a protein. The color shows the intensity of the respective protein and missing values are greyed out. The samples and proteins are clustered using a hierarchical clustering on the expected distances calculated with Equation (42). The annotations on top of the heatmap indicate which samples were compared in the different performance and calibration experiments shown in Figure 4 and Supplementary Figure S6-S8.

**Suppl. Figure S3:**
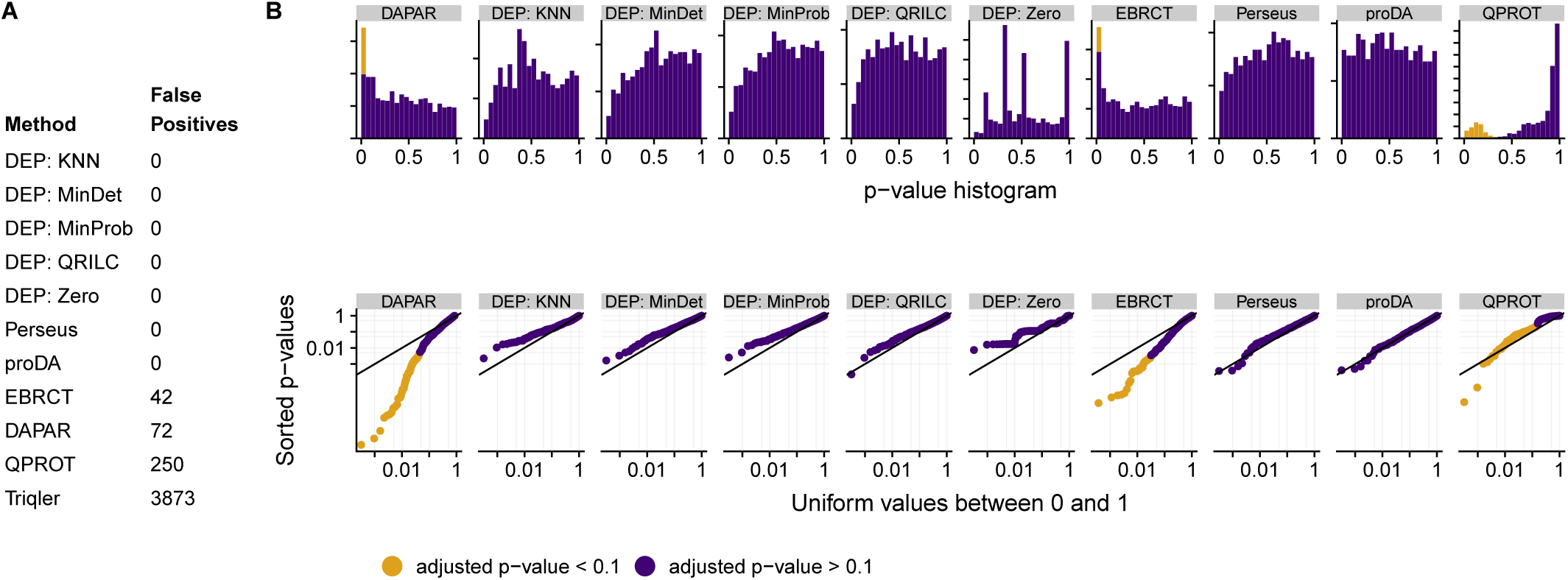
Same as Supplementary Figure 3, but for a comparison of 4 vs 4 samples

**Suppl. Figure S4:**
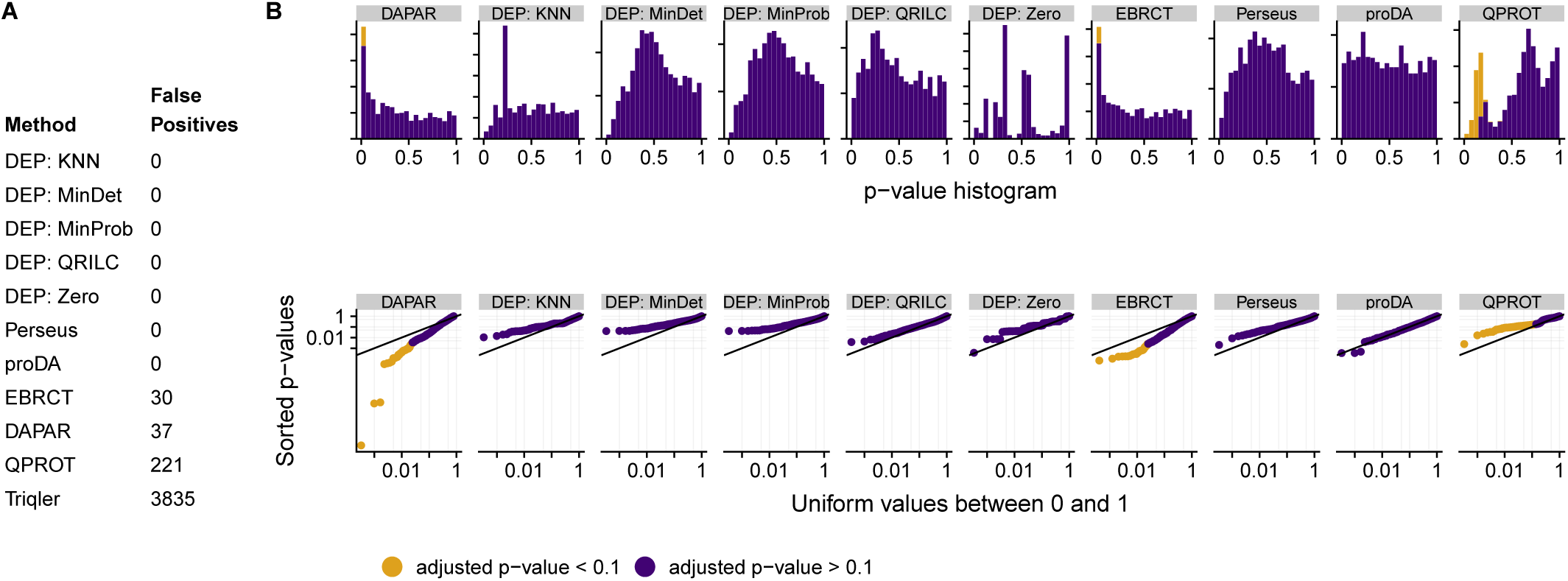
Same as Supplementary Figure 3, but for a comparison of 4 vs 4 samples

**Suppl. Figure S5:**
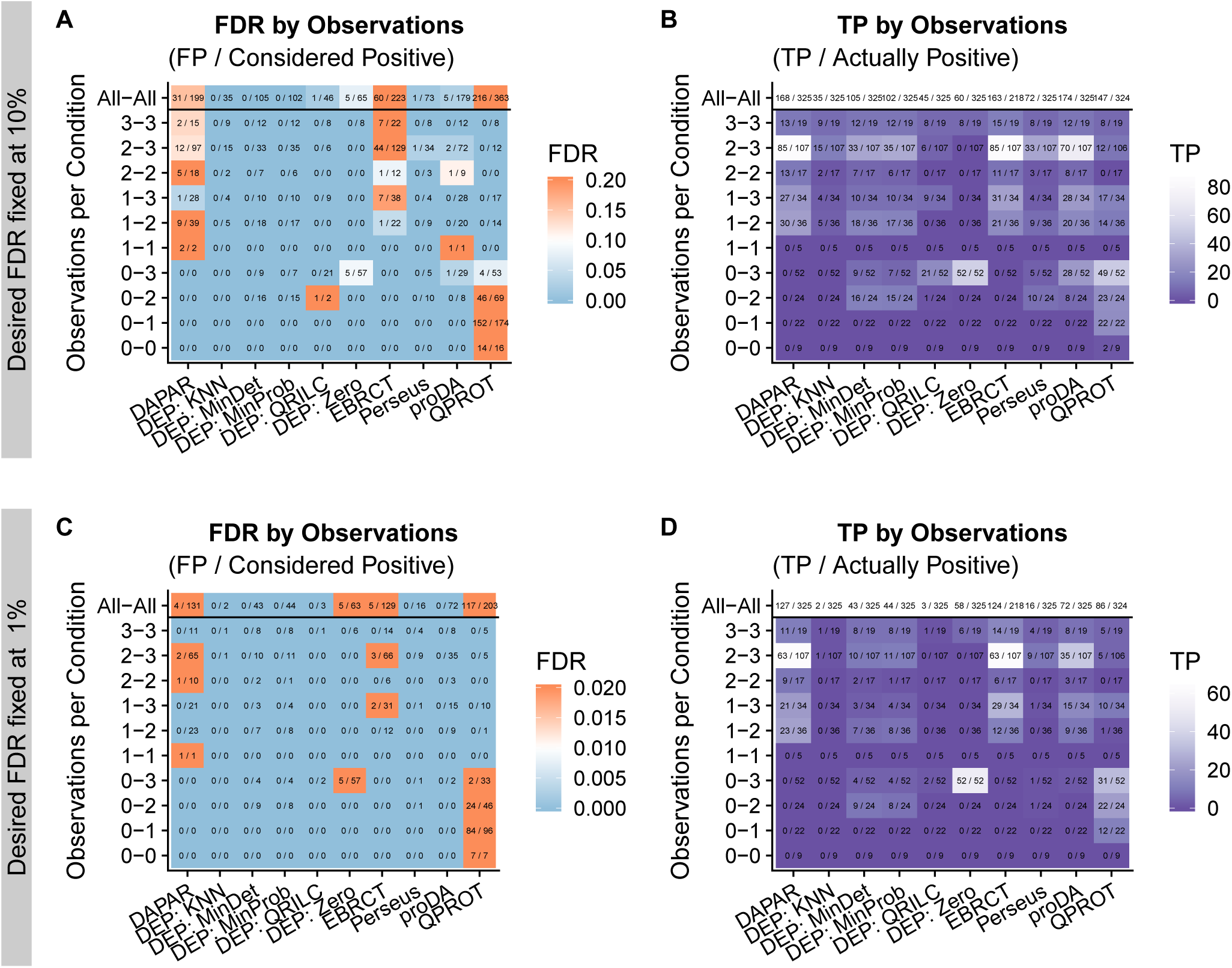
Calibration and performance stratified by the number of observations on the three vs three comparison on the de Graaf dataset with 20% changed proteins. The x-axis shows the six tested methods and the y-axis the number of observed values in condition one and two. The smaller number is listed first. In addition the top line on the y-axis shows the marginal over all combination of observations. A) and B) show the results when fixing the desired FDR at 10%. A) shows the actual FDR for each method and B) shows the number true positives. and D) show the same features if the desired FDR is fixed at 1%. The color scales show the FDR (A and C) or the number of true positives (B and D). White indicates the optimal value. In A) and C) light blue color indicates a conservative FDR, whereas orange indicates an anti-conservative FDR.

**Suppl. Figure S6:**
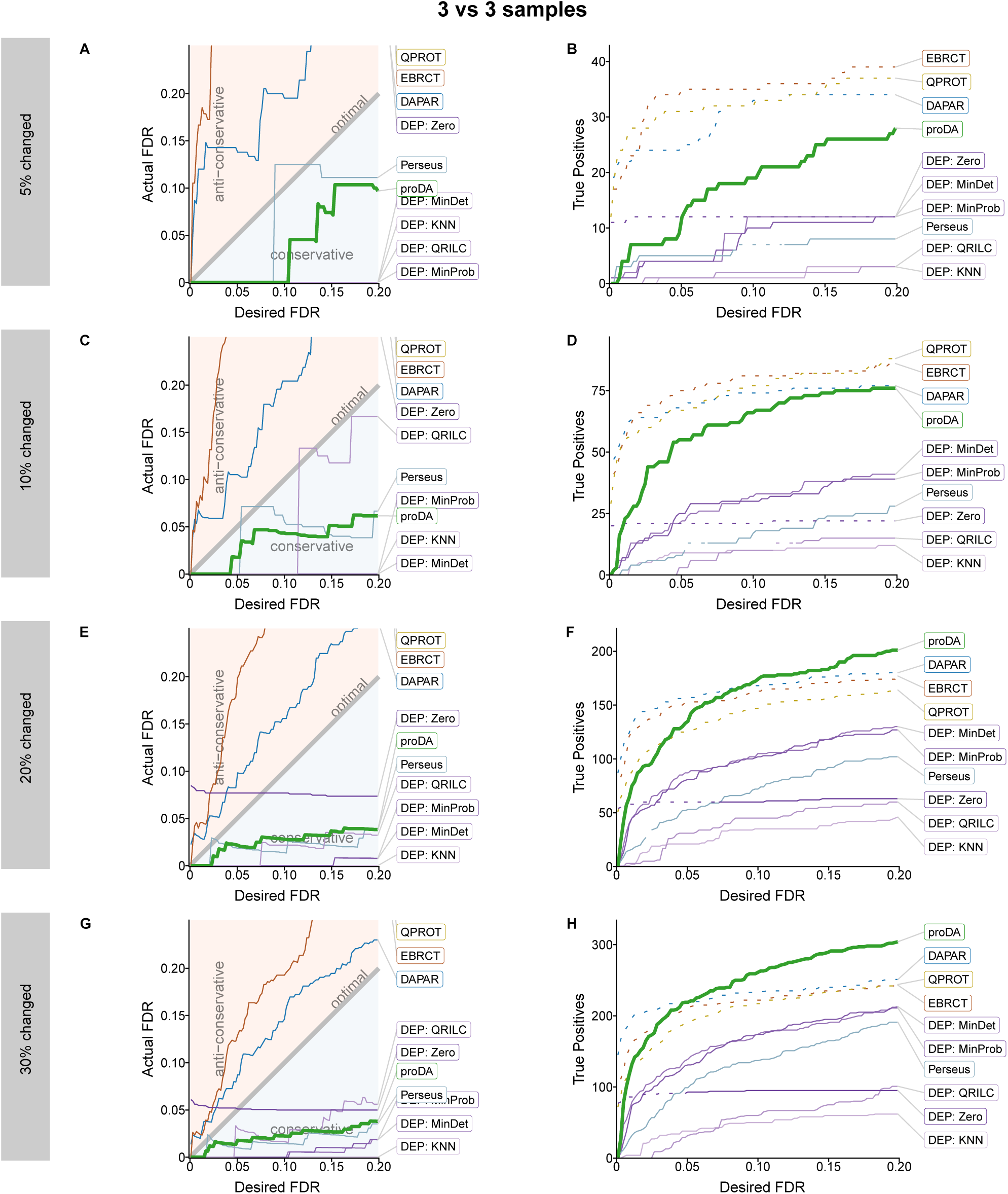
Calibration and performance comparison with three vs. three samples. A,C,E,G) comparison of the desired FDR with the FDR that is actually produced by the tool acording to the ground truth. The line for the QPROT method is missing because it is literally of the charts. B,D,F,H) Plot of how many actually changed proteins (true positives) each method identified at a specified FDR level. The regions where a method failed to control the FDR in the left column are plotted as dashed lines. Note that for the performance comparison with only 5% of the proteins changed, in absolute terms there are only a handful of false positives and the corresponding estimates of the FDR are very noisy.

**Suppl. Figure S7:**
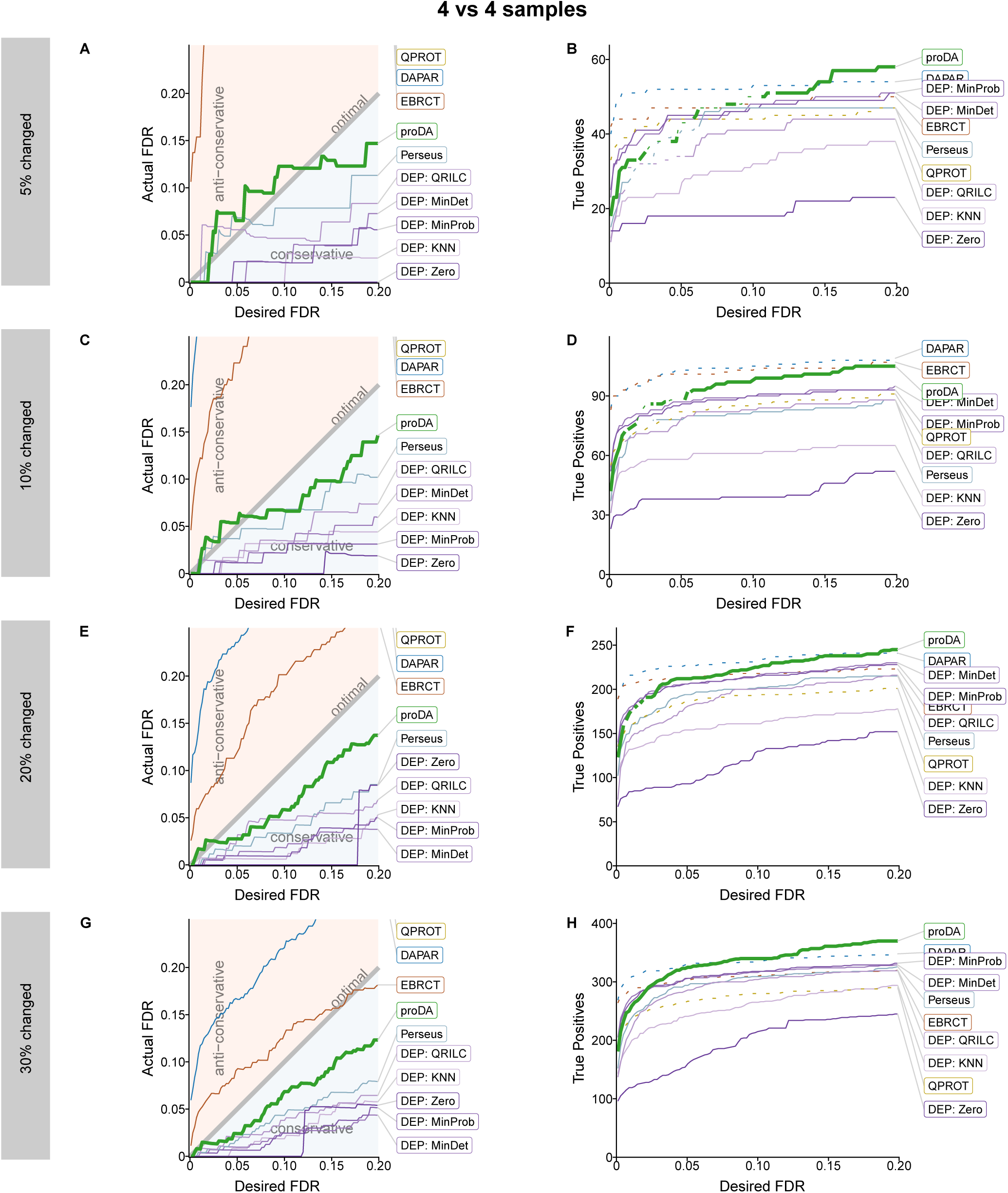
Same as Supplementary Figure S6, but for a comparison of 4 vs 4 samples.

**Suppl. Figure S8:**
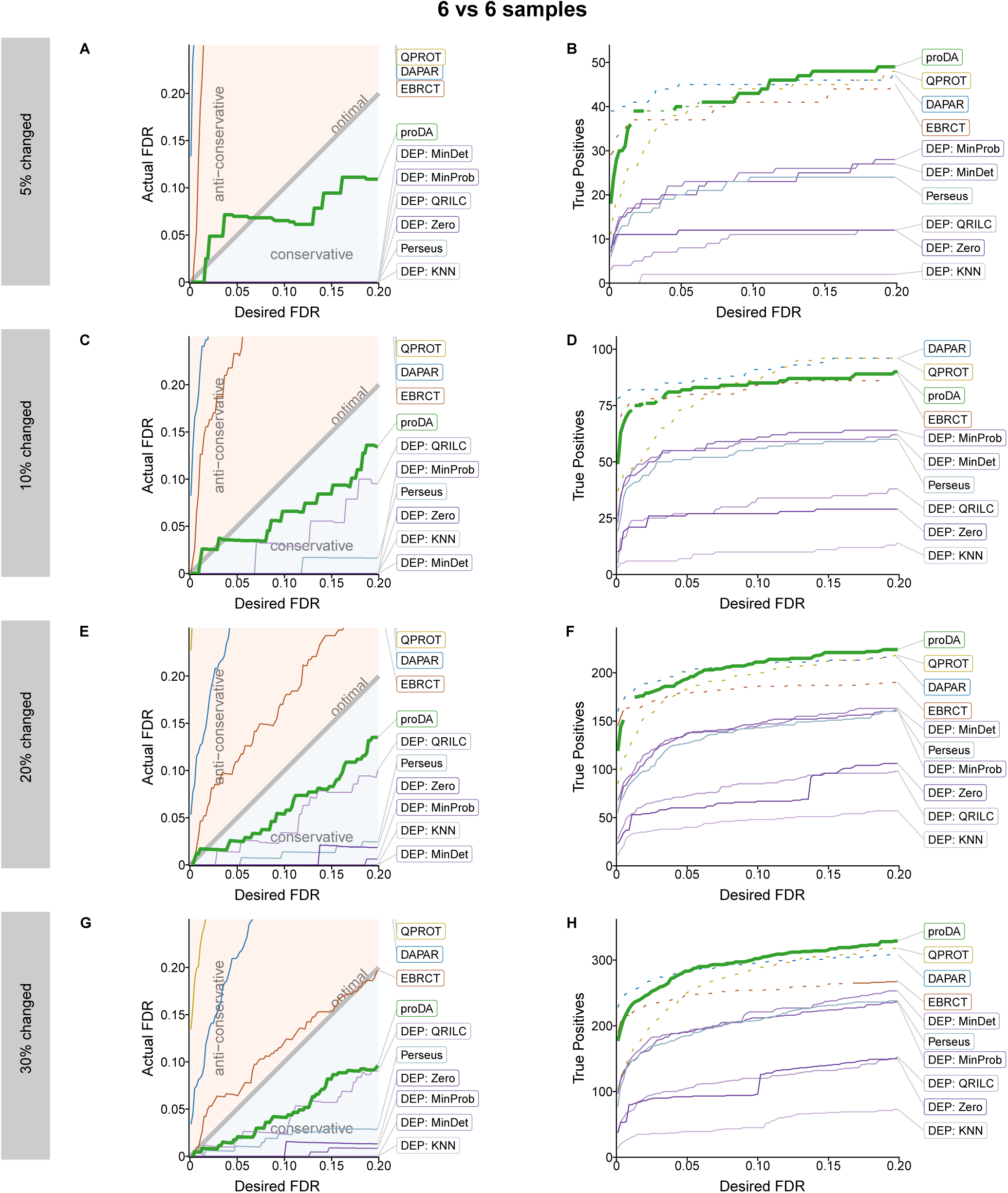
Same as Supplementary Figure S6, but for a comparison of 6 vs 6 samples.

